## Supplementary Material for "Predicting the diversity of photosynthetic light-harvesting using thermodynamics and machine learning"

### 1 Pigment lineshapes

| Pigment (abbreviation)$^{origin}$ | Absorption peaks (nm) | Peak $\sigma$ (nm) | Amplitudes |
|---|---|---|---|
| Bacteriochlorophyll *b* (BChl *b*)[1] | 805, 790, 730, 685, 615 | 15.0, 25.0, 20.0, 15.0, 17.0 | 0.933, 0.16, 0.15, 0.14, 0.3 |
| Bacteriochlorophyll a (BChl a)[1] | 780, 728, 612 | 13.5, 34.6, 18.4 | 0.933, 0.16, 0.230 |
| Chlorophyll f (Chl f)[1] | 707, 679, 650 | 11.0, 18.0, 30.0 | 0.75, 0.18, 0.08 |
| Chlorophyll d (Chl d)[1] | 698, 665, 650 | 9.0, 10.0, 30.0 | 0.75, 0.06, 0.1 |
| Chlorophyll a (Chl a)[1] | 671, 641, 617, 615 | 8.5, 10.5, 7.0, 30.0 | 0.87, 0.13, 0.05, 0.1 |
| Chlorophyll b (Chl b)[1] | 654, 623, 603, 562 | 10.0, 14.0, 11.0, 30.0 | 0.91, 0.12, 0.16, 0.15 |
| Allophycocyanin (APC)[2] | 655, 625, 595 | 10.0, 25.0, 50.0 | 0.650, 0.66, 0.35 |
| Phycocyanin (PC)[2] | 625, 608, 585 | 11.0, 30.0, 45.0 | 0.750, 0.220, 0.95 |
| phycoerythrin (PE)[2] | 565, 545, 520 | 11.5, 20.0, 25.0 | 0.50, 0.40, 0.220 |
| r-phycoerythrin (r-PE)[3] | 568, 542, 496 | 9.0, 25.0, 10.0 | 0.60, 1.080, 0.680 |

Table 1: Table of fit information (pigment name, absorption peaks, widths and amplitudes) for absorption spectra for use in the antenna model and genetic algorithm. Subscripts denote origin of experimental spectrum: (1) PhotoChemCAD [1], (2) [2], (3) [3]. All lineshapes taken from isolated pigments in solution. Fits to experimental data shown below in Figs. 1 to 3.

| Pigment (abbreviation)$^{origin}$ | Emission peaks (nm) | Peak $\sigma$ (nm) | Amplitudes |
|---|---|---|---|
| Bacteriochlorophyll *b* (BChl *b*)[1] | 821, 840 | 13.5, 25.0 | 0.933, 0.16 |
| Bacteriochlorophyll a (BChl a)[1] | 793, 810 | 11.0, 40.0 | 0.9, 0.15 |
| Chlorophyll f (Chl f)[1] | 722, 740 | 13.0, 40.0 | 0.85, 0.11 |
| Chlorophyll d (Chl d)[1] | 705, 730, 765 | 10.5, 11.0, 20.0 | 0.85, 0.06, 0.07 |
| Chlorophyll a (Chl a)[1] | 677, 695, 732 | 9.0, 11.0, 20.0 | 0.85, 0.10, 0.11 |
| Chlorophyll b (Chl b)[1] | 662, 688, 720 | 9.5, 11.0, 18.0 | 0.85, 0.09, 0.16 |
| Allophycocyanin (APC)[2] | 662 | 16.5 | 1.0 |
| Phycocyanin (PC)[2] | 600, 645, 690 | 11.0, 18.0, 15.0 | 0.10, 0.850, 0.05 |
| phycoerythrin (PE)[2] | 585, 600 | 10.5, 25.0 | 0.85, 0.20 |
| r-phycoerythrin (r-PE)[3] | 578, 600, 630 | 10.0, 30.0, 20.0 | 0.90, 0.35, 0.12 |

Table 2: Table of fit information (pigment name, absorption peaks, widths and amplitudes) for emission spectra for use in the antenna model and genetic algorithm. Subscripts denote origin of experimental spectrum: (1) PhotoChemCAD [1], (2) [2], (3) [3]. All lineshapes taken from isolated pigments in solution. Fits to experimental data shown below in Figs. 1 to 3.

We use a range of pigment lineshapes absorbing light of different wavelengths which are taken from various organisms in different photosynthetic niches. For each subunit of the antenna the algorithm is free to choose any of the available pigment types when constructing or mutating the antennae.

This choice is implemented by taking published absorption and emission lineshapes for each pigment type and fitting them to a set of Gaussians. The available choices along with the fitted Gaussian parameters are given in Tables 1 and 2, as well as in the file `pigments/pigment_data.json` available on github. These fitted Gaussians are then recalculated for the set of wavelengths given in the input spectra to avoid any interpolation issues and also to allow for future work where it may be necessary to introduce shifting of absorption peaks.

The absorption and emission spectrum for each pigment type is shown in Figs. 1 to 3.

### 2 Antenna compositions for sweep over cost parameter

The following figures show the averaged absorption spectra and pigment compositions of the inner 10 subunits for each set of incident light conditions as a function of $\chi$. Colour coding is consistent throughout and shown as a legend on each subfigure.

Fig. 4 shows antenna compositions for full sunlight as a function of intensity and cost. We see a mix of APC and PC mostly, with a switch to Chl *a* at lower intensity and higher cost, and a general increase in the size of the antennae as the cost gets lower.

Figs. 5 to 7 show the antenna compositions for red light (top rows), shallow water (middle rows) and deep water (bottom rows) as a function of light intensity and cost.

#### 2.1 Electron output and efficiency of antennae

Fig. 8 shows the electron outputs and efficiencies for each light environment at each of the simulated intensities as a function of $\chi$.

### 3 Far-red light illumination

The case of far-red light illumination is worth exploring in more detail. In Fig. 4 of the main text we present results using decreasing intensities of visible light below 700nm with a reaction centre absorbing at 720nm. Fig. 9 shows antenna composition histograms for the results presented in the main text for high $\chi = 0.03$, medium $\chi = 0.02$ and low $\chi = 0.005$ cost values, showing the switch from Chls $a$ and $b$ to $d$ at lower intensities and finally Chl $f$ at the lowest intensities and highest cost value.

### 4 Effect of contingent evolution in deep water as a function of cost

Antenna composition histograms are shown in Fig. 10 which correspond to the absorption spectra shown in main text Fig. 5. We see that our *in-situ* evolved antennae are substantially enriched with PE compared to their pre-adapted counterparts for each cost value; the pre-adapted population retains a core of either APC or Chl $b$, which have very similar absorption profiles, and hence their absorption has a much larger peak at around 650nm compared to the *in-situ* population. Regardless, however, we see that the *in-situ* population is composed of broadly the same sets of pigments, just in differing ratios.

The electron outputs and efficiencies are very similar for the two populations as mentioned in the main text; details are shown in Fig. 11.

### 5 Antenna composition for anoxygenic and oxygenic antennae for different stellar temperatures

Figs. 12 to 14 show antenna composition histograms for Fig. 6 of the main text for costs $\chi = 0.03, 0.02, 0.005$ respectively. We see predominantly BChl $b$ for the anoxygenic reaction centre for all stellar temperatures and cost values, with more BChl $a$ and Chls $f$ and $d$ mixed in as the stellar temperature increases and cost decreases.

For oxygenic we find Chl $d$ antennae for M-dwarf illumination at high cost, giving way to predominantly Chl $a$ and $b$ as well as APC and PC at higher stellar temperatures. We also note that under M-dwarf illumination the anoxygenic antennae are much larger than oxygenic, since there is comparatively much more available light for photosynthesis; this is reflected in the significantly higher electron output discussed in the main text.

#### 6 Code

The code used in this work will be available at `https://github.com/QMUL-DuffyLab/gala` upon publication.

Both Python and Fortran versions of the non-negative least squares solver used to calculate electron output and efficiency for a given antenna structure are included, but only the Python version is needed to run simulations. The only requirements then are a modern version of Python ($\geq 3.9$) and various standard packages (numpy, scipy, matplotlib), all of which are available via `pip` and/or `conda`. The included README together with docstrings in the code should be sufficient to set up and run simulations; please with any questions.

#### 7 Additional data

A complete set of the output data used in this work is available at `https://doi.org/10.5281/zenodo.14514090`, including additional sets of simulations for different values of the cost parameter for various light environments and additional visible light intensities for far red light.

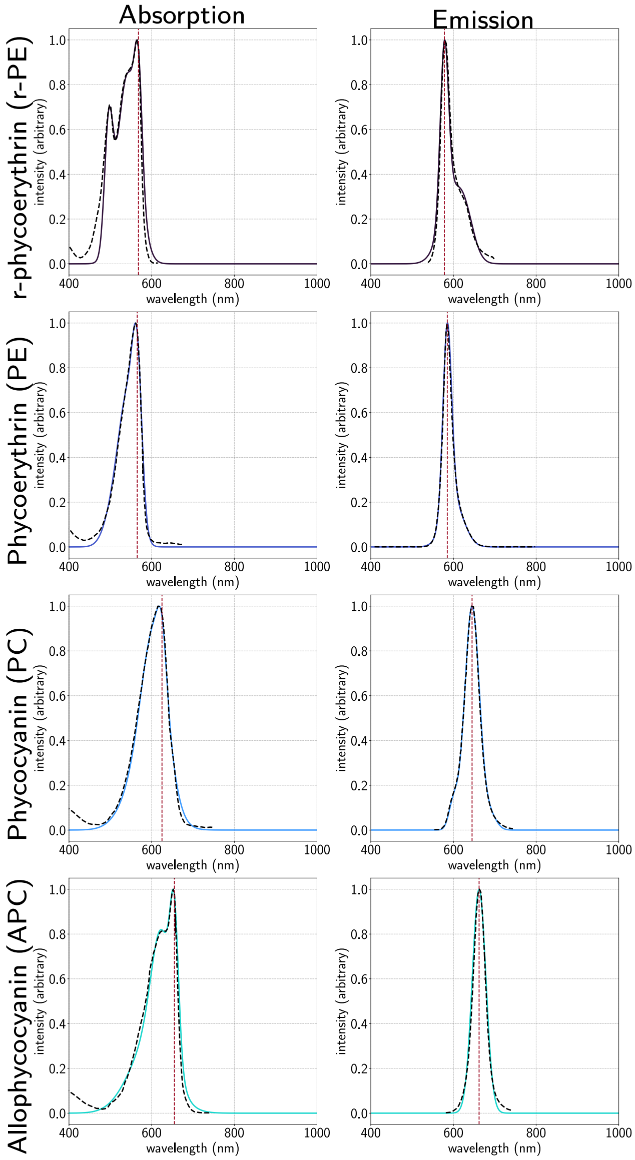


Figure 1: Fits to absorption and emission lineshapes for r-phycoerythrin, phycoerythrin, phycocyanin and allocphycocyanin. Fits shown as solid lines, experimental data as dashed lines. The vertical dashed lines on each plot denote the 0-0 line used for determining energy gaps between pairs of pigments.

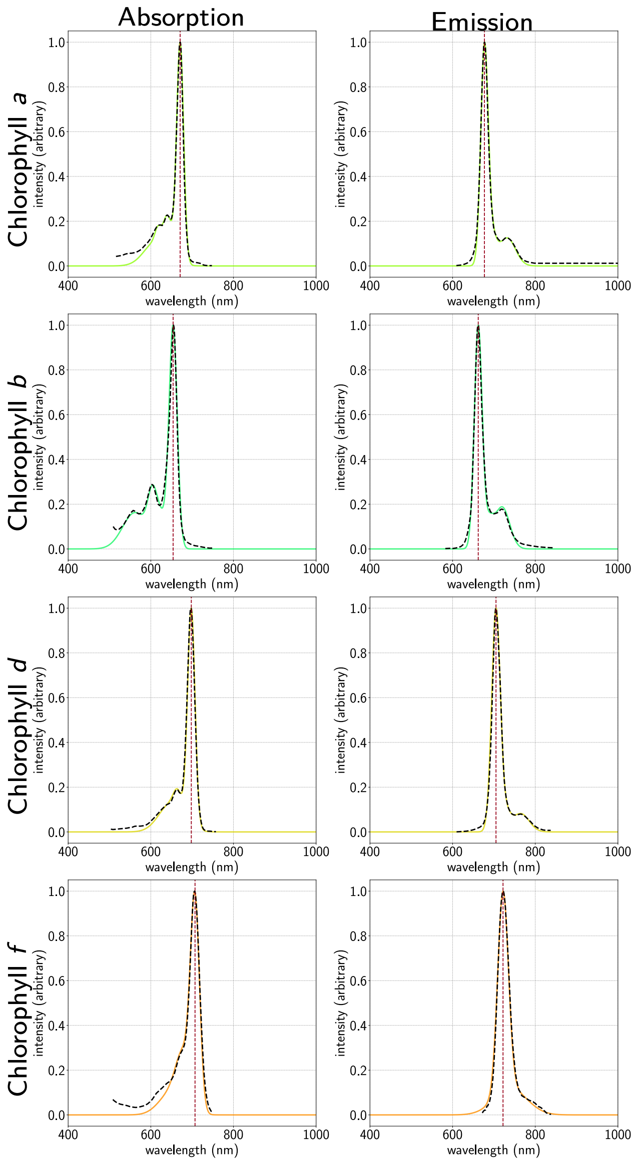


Figure 2: Fits to absorption and emission lineshapes for Chl *a*, Chl *b*, Chl *d* and Chl *f*. Fits shown as solid lines, experimental data as dashed lines. The vertical dashed lines on each plot denote the 0-0 line used for determining energy gaps between pairs of pigments.

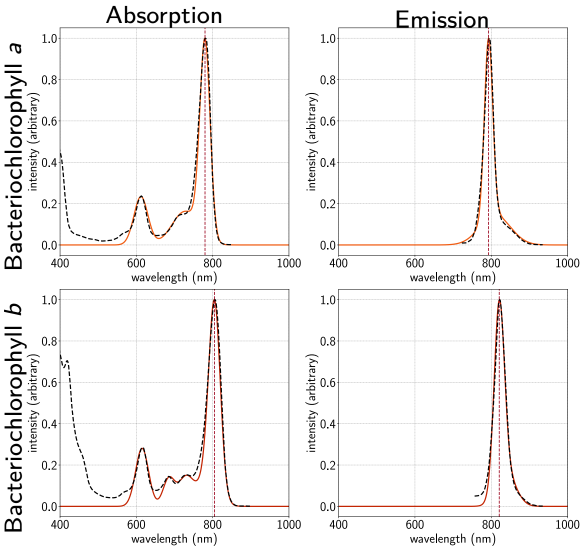


Figure 3: Fits to absorption and emission lineshapes for BChl *a* and *b*. Fits shown as solid lines, experimental data as dashed lines. The vertical dashed lines on each plot denote the 0-0 line used for determining energy gaps between pairs of pigments.

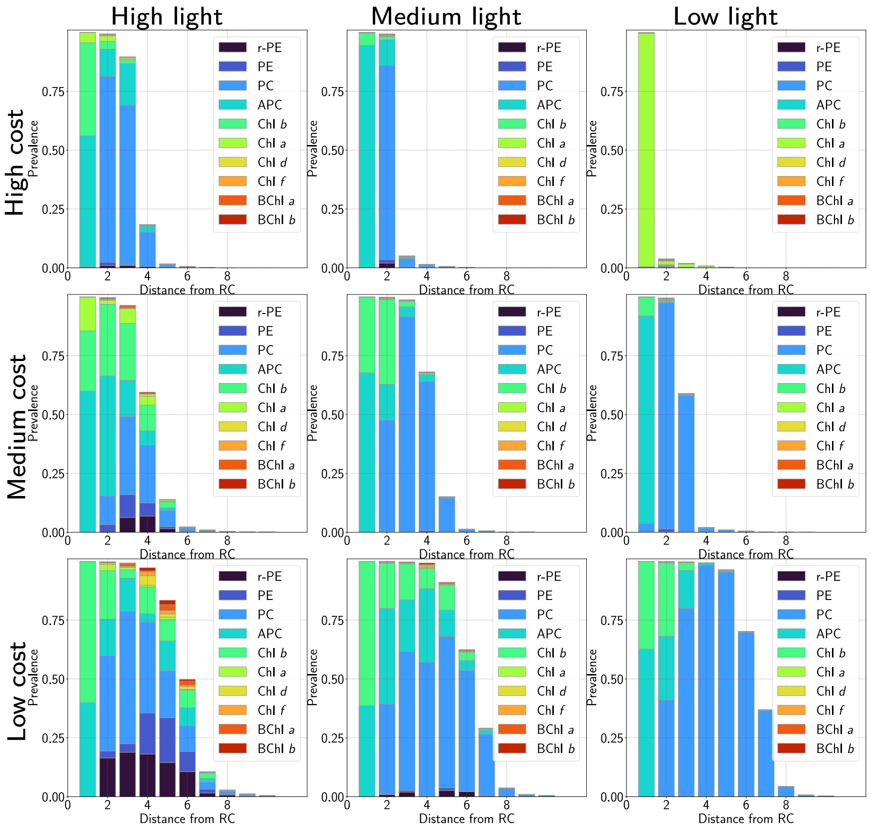


Figure 4: Absorption spectra (left) and pigment composition (right) of antenna evolved under AM1.5 full sunlight illumination as a function of light intensity and cost $\chi$. Columns correspond to differing intensities: full (left column), medium (300 $\mu$E) (middle column) and low (50 $\mu$E) (right column). Rows correspond to differing cost values: high ($\chi = 0.04$) (top row), medium ($\chi = 0.02$) (middle row) and low ($\chi = 0.005$) (bottom row).

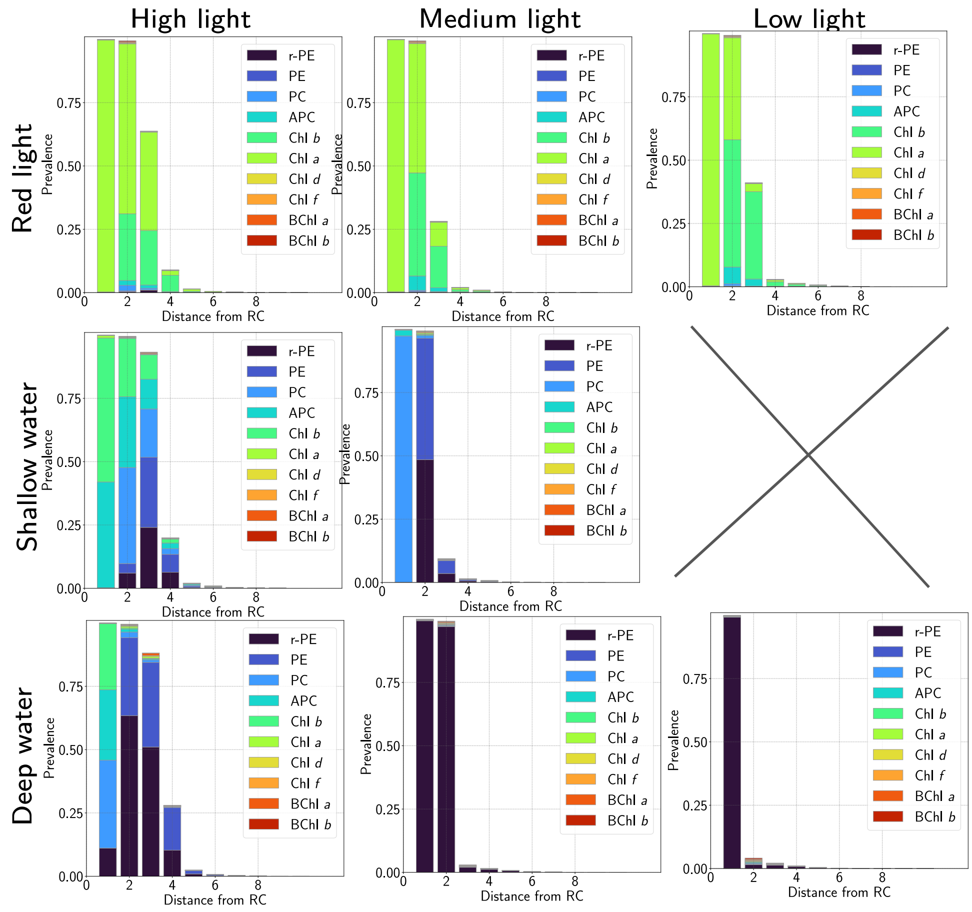


Figure 5: Antenna composition histograms for the highest cost value simulated: $\chi = 0.04$ for high and medium intensity (left and middle columns), $\chi = 0.01$ for low intensity (right column). Rows correspond to differing illumination: (top row) red light, (middle row) shallow water and (bottom row) deeper water, as in the main text. Note that at low intensity in shallow water with cost $= \chi = 0.01$ the algorithm fails to find any viable antennae; as mentioned in the main text, we suggest that this is due to the algorithm becoming increasingly sensitive to small features in the input spectrum as the light intensity decreases.

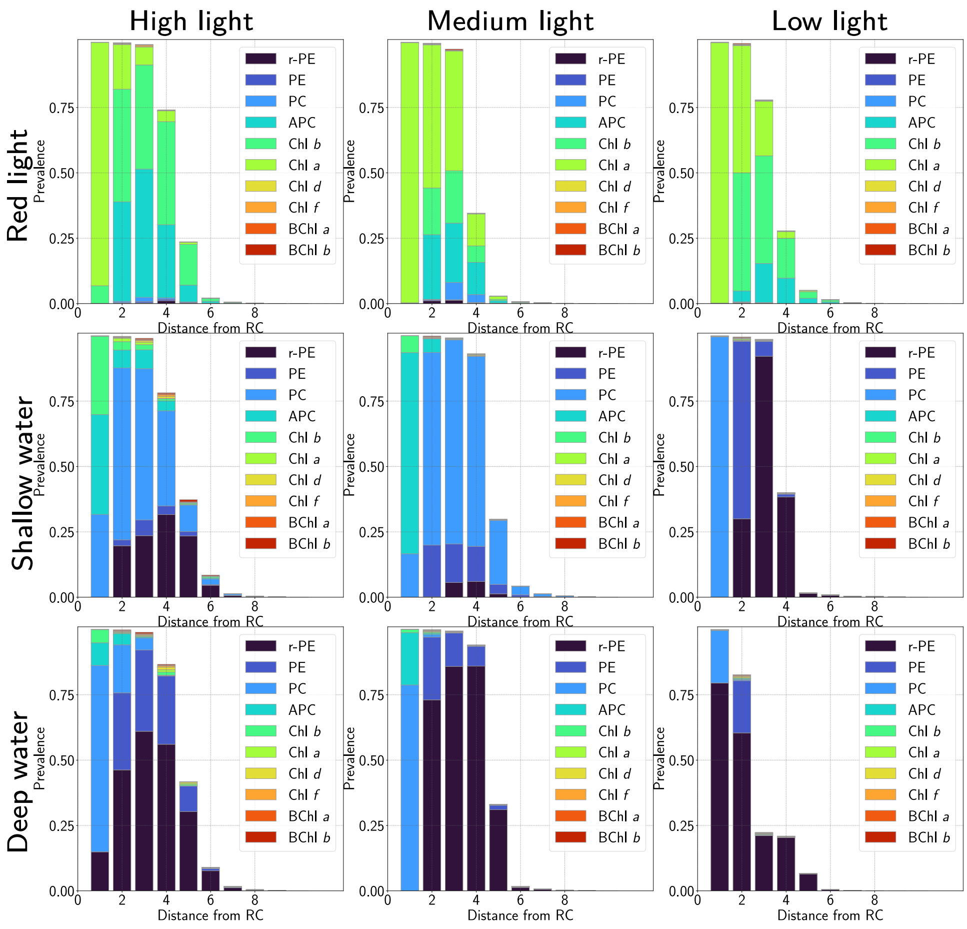


Figure 6: Antenna composition histograms for the medium cost value simulated: $\chi = 0.02$ for high and medium intensity (left and middle columns), $\chi = 0.008$ for low intensity (right column). Rows correspond to differing illumination: (top row) red light, (middle row) shallow water and (bottom row) deeper water, as in the main text.

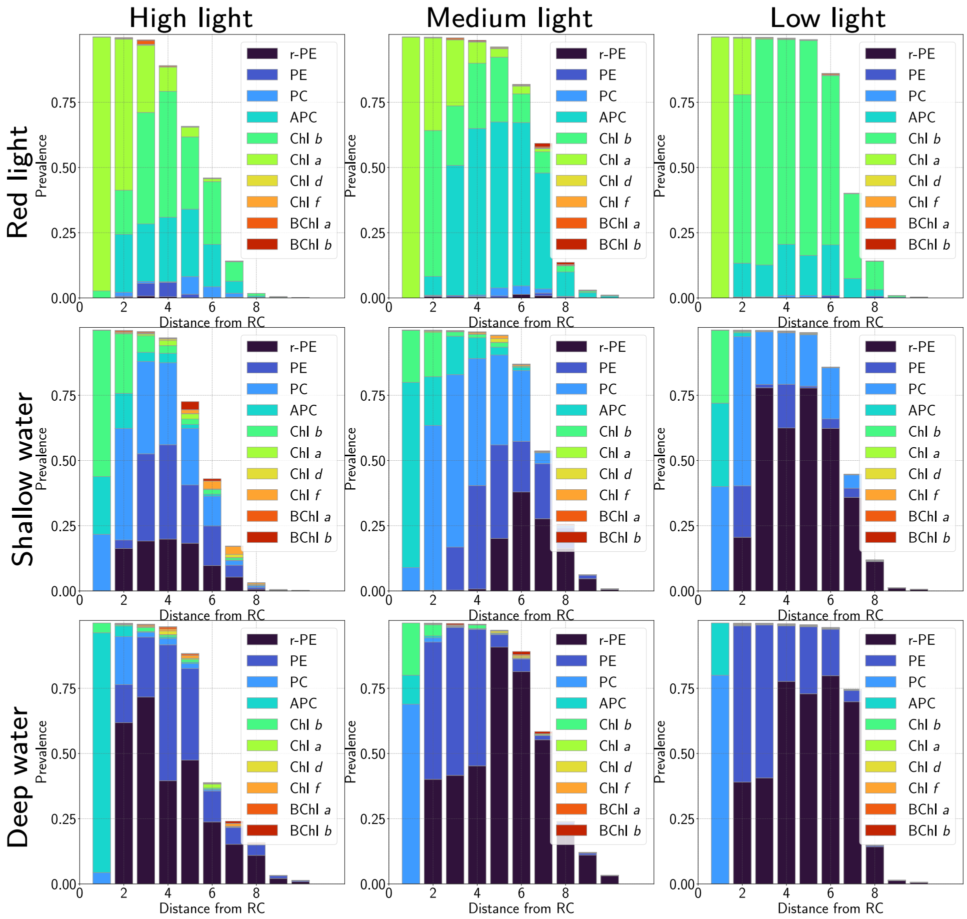


Figure 7: Antenna composition composition histograms for the lowest cost value simulated: $\chi = 0.005$ for high and medium intensity (left and middle columns), $\chi = 0.004$ for low intensity (right column). Rows correspond to differing illumination: (top row) red light, (middle row) shallow water and (bottom row) deeper water, as in the main text.

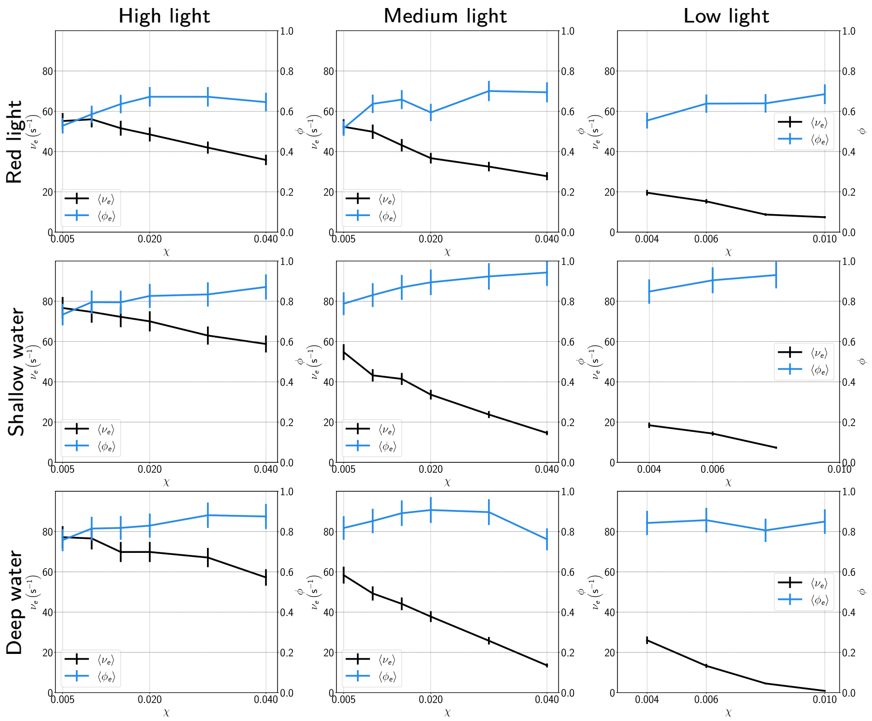


Figure 8: Electron output $\nu_e$ (black lines) and efficiency $\phi$ (blue lines) for differing light environments (red light, top row; shallow water, middle row; deep water; bottom row) and intensities (high, left column; medium, middle column; low, right column) as in the main text, as a function of cost $\chi$.

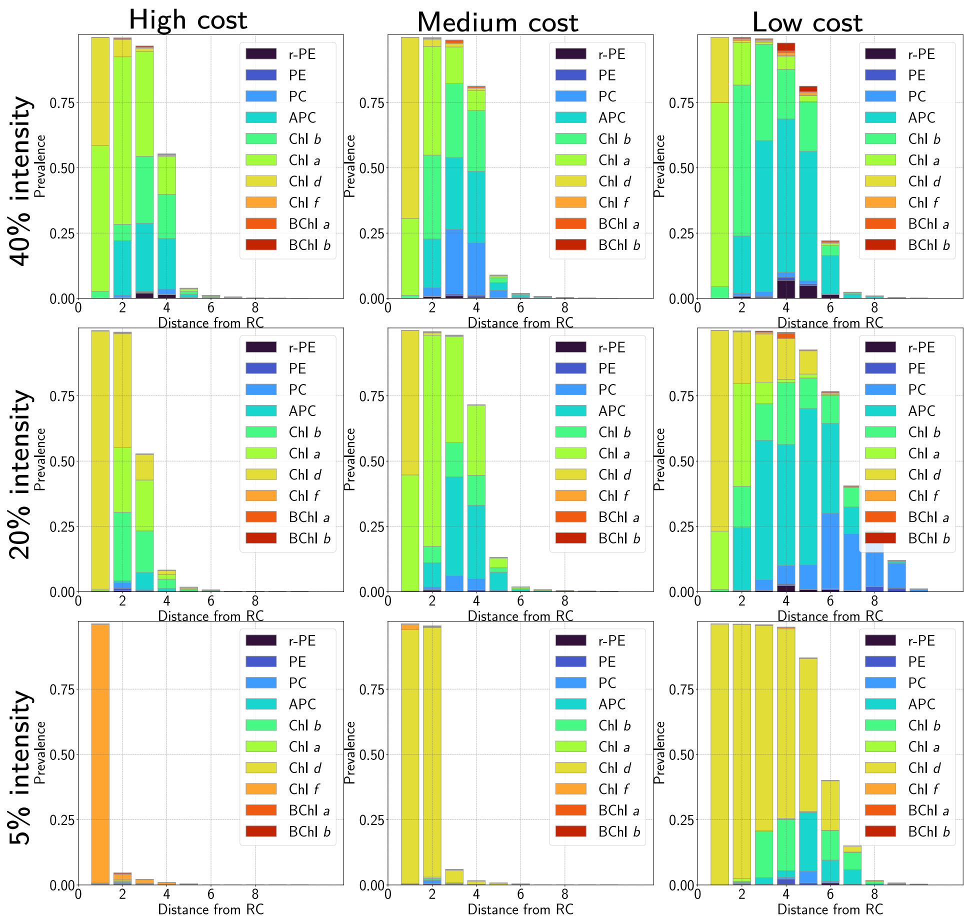


Figure 9: Antenna composition histograms for different cost values and visible light intensities with a far-red adapted reaction centre. Columns: high cost (left column) $\chi = 0.03$, medium (middle column) $\chi = 0.02$, low (right column) $\chi = 0.005$. Rows correspond to differing visible light intensities: (top row) 40% of visible light compared to full sunlight, (middle row) 20% and (bottom row) 5%, as in the main text. We see a change from Chls $a$ and $b$, with some APC and PC included for lower cost values, to mostly Chl $d$ antennae as the light intensity decreases, and finally a small Chl $f$ antenna at high cost and low intensity.

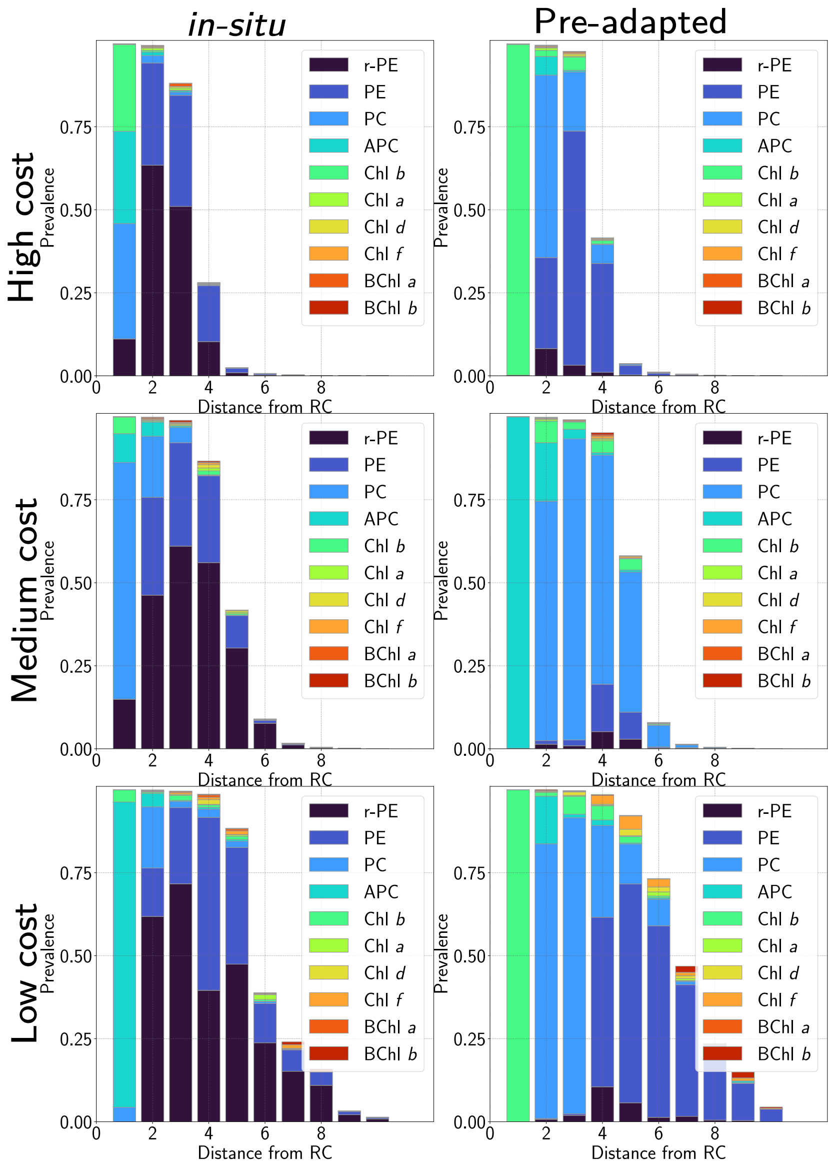


Figure 10: Antenna composition histograms for different cost values for *in-situ* evolved antennae (left column) and pre-adapted antennae (right column).. Rows correspond to different cost values: (top row) $\chi = 0.04$ (middle row) $\chi = 0.02$ and (bottom row) $\chi = 0.005$ as in the main text. We see that the *in-situ* evolved population contains signficantly more phycoerythrin for all cost values, and substantially less APC and Chl $b$.

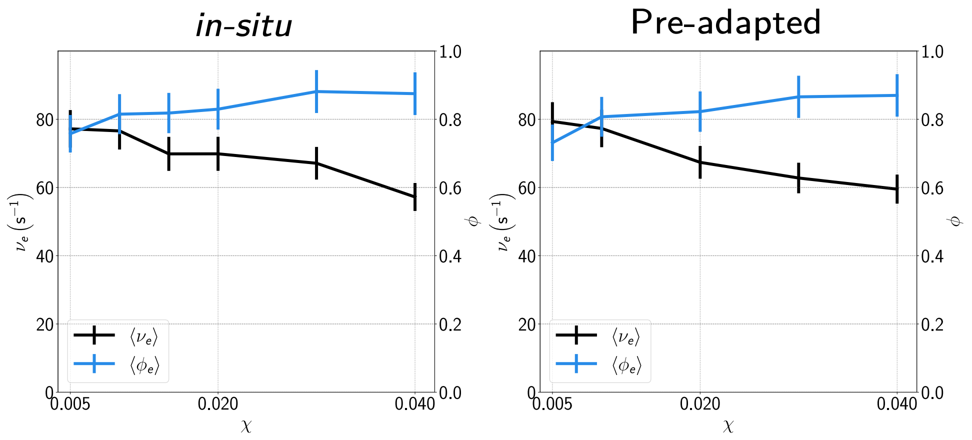


Figure 11: Electron outputs and antenna efficiencies as a function of cost for *in-situ* and pre-adapted populations, showing very similar antenna performance between the two populations. Standard error for each value shown as vertical bars.

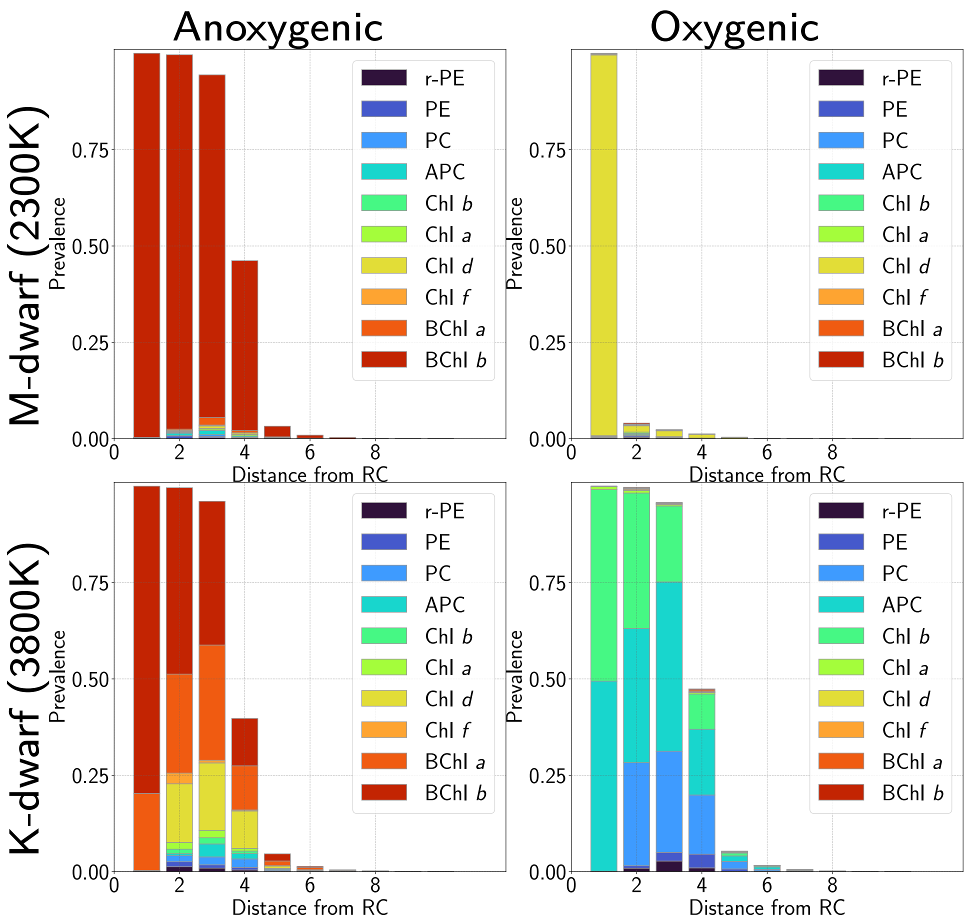


Figure 12: Antenna composition histograms for M-dwarf (2300K, top row) and K-dwarf (3800K, bottom row) for anoxygenic (left column) and oxygenic (right column) reaction centres, with high cost $\chi = 0.03$.

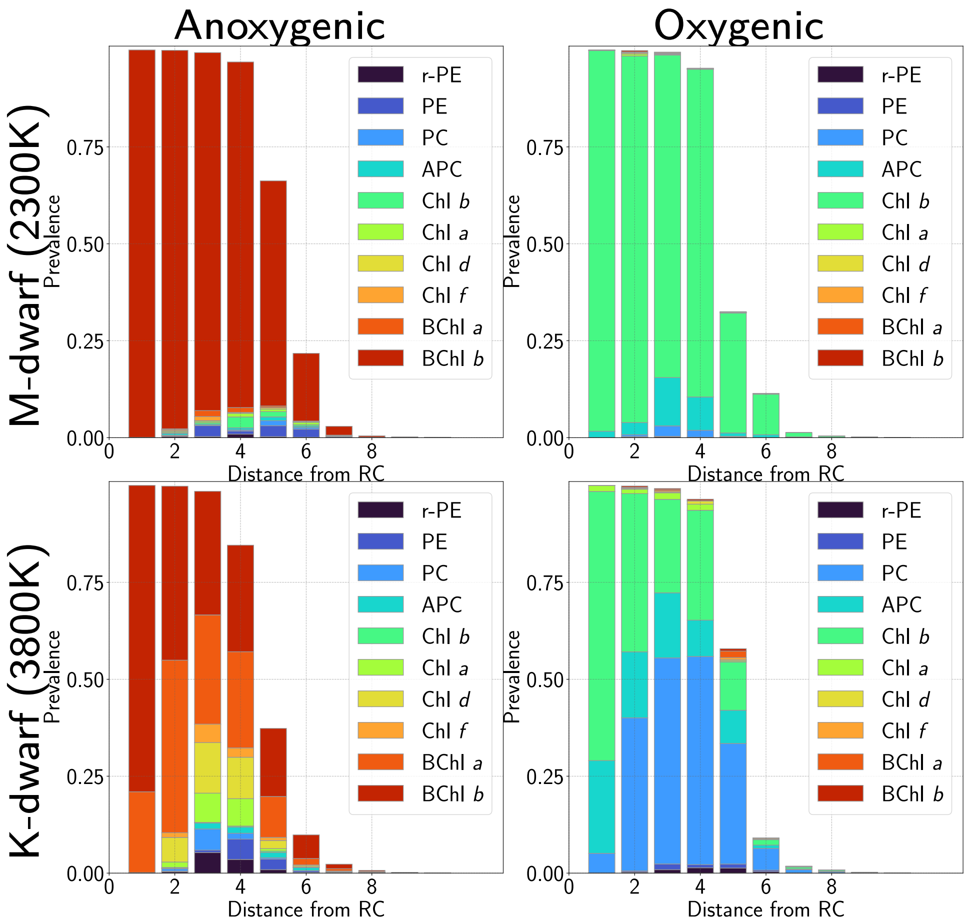


Figure 13: Antenna composition histograms for M-dwarf (2300K, top row) and K-dwarf (3800K, bottom row) for anoxygenic (left column) and oxygenic (right column) reaction centres, with medium cost $\chi = 0.02$.

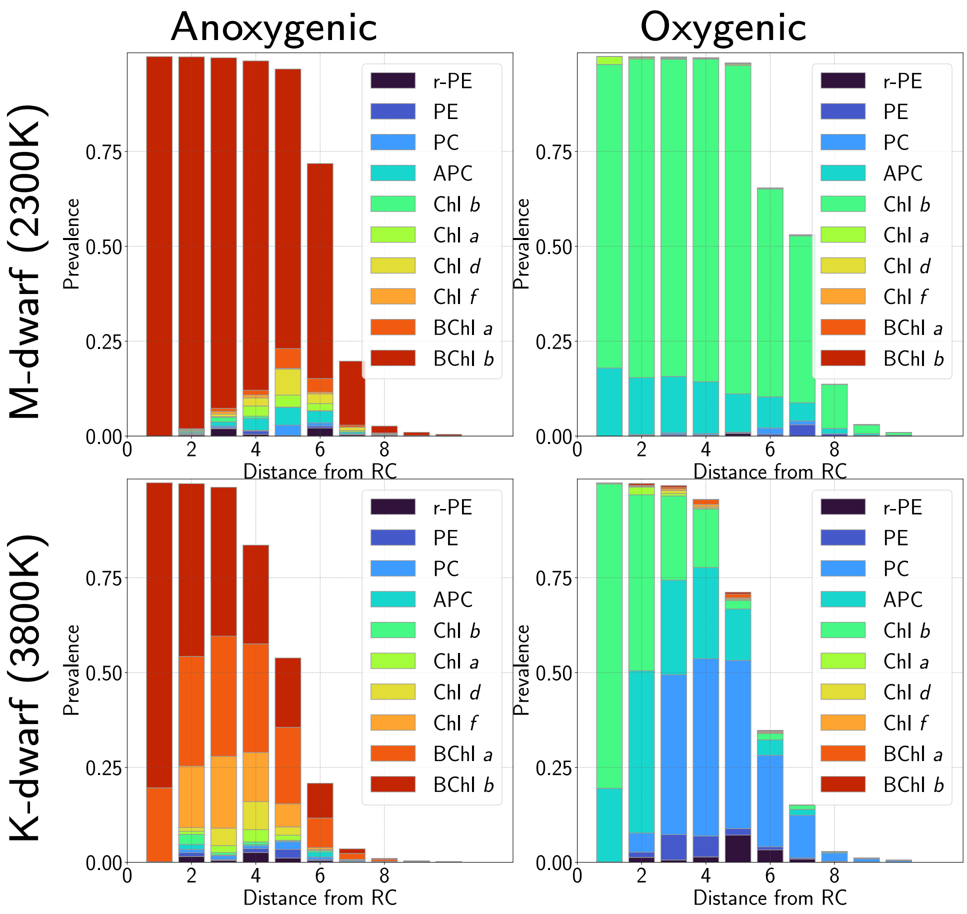


Figure 14: Antenna composition histograms for M-dwarf (2300K, top row) and K-dwarf (3800K, bottom row) for anoxygenic (left column) and oxygenic (right column) reaction centres, with low cost $\chi = 0.005$.
